# Differential nanoscale organisation of LFA-1 modulates T cell migration

**DOI:** 10.1101/602326

**Authors:** Michael J. Shannon, Judith Pineau, Juliette Griffié, Jesse Aaron, Tamlyn Peel, David J. Williamson, Rose Zamoyska, Andrew P. Cope, Georgina H. Cornish, Dylan M. Owen

## Abstract

Effector T-cells rely on integrins to drive adhesion and migration to facilitate their immune function. Heterodimeric transmembrane integrin LFA-1 (αLβ2) regulates adhesion and migration through linkage of the extracellular matrix with the intracellular actin treadmill machinery. We quantitated the velocity and direction of F-actin flow in migrating T-cells alongside single molecule localisation of transmembrane and intracellular LFA-1. Our results show that retrograde actin flow positively correlated and immobile actin negatively correlated with T-cell velocity. Plasma membrane localised LFA-1 forms unique nano-clustering patterns in the leading edge, compared to the mid-focal zone, in migrating T-cells. Deleting the cytosolic phosphatase PTPN22, a negative regulator of integrin signaling, increased T-cell velocity, and leading-edge cluster co-localisation of pY397 FAK, pY416 Src family kinases and LFA-1. These data suggest that differential nanoclustering patterns of LFA-1 in migrating T-cells can instruct intracellular signalling linked with the actin treadmill. Our data presents a paradigm where T cells modulate the nanoscale organisation of adhesion and signalling molecules to fine tune their migration speed. This has implications for the regulation of immune and inflammatory responses.

## Introduction

Integrin lymphocyte functional antigen-1 (LFA-1; αLβ2) is a transmembrane heterodimer highly expressed on T-cells. It is essential for T-cell function and is required for naïve lymphocyte antigenic activation, recruitment of immune cells to sites of infection and cytotoxic killing. LFA-1 expressed on effector T cells binds to intracellular adhesion molecule (ICAM)-1 to induce rapid migration *in vivo* and *in vitro* (Hons et al. 2018; Teijeira et al. 2017). The force required for migration is generated by actin polymerisation and myosin driven actin flow, and integrin adhesion molecules on the cell surface which couple the actin flow to the underlying substrate (Shannon et al. 2015). Actin and integrins are indirectly linked via a group of proteins collectively known as the molecular clutch, which regulates this mechanical coupling (Ishibashi et al. 2015; Chen et al. 2012; Case & Waterman 2015). The control of this linkage is integral to achieving high-speed migration (Hons et al. 2018), and is especially important during inflammation (Teijeira et al. 2017). It has mostly been investigated in terms of integrin affinity, where low, intermediate and high affinity states describe a specific pattern of localisation within a polarised migrating T cell that correlates to adhesion strength. Low and intermediate affinity integrin are located in the leading edge, whereas high affinity LFA-1 is located in the mid cell body – the focal zone (Smith et al. 2005).

Control of adhesion is also achieved through actin linkers (talin, vinculin) and regulatory kinases (FAK, Src family kinases)(Nordenfelt et al. 2016; Raab et al. 2017a), common to other migrating cell types, that form large micron sized focal adhesions(Kanchanawong et al. 2010). In T cells, these complexes are much smaller (nanoscale), containing only a few molecules (Burn et al. 2016) but similar in size and composition to nascent adhesions (Changede et al. 2015). Their spatial organisation therefore represents a novel paradigm for nanoscale regulation which to date has not been studied in as much depth.

Conventional fluorescence microscopy is limited in resolution to around 200 nm. Recently, a number of super-resolution approaches have been developed which overcome this limitation. Single molecule localisation microscopy (SMLM), here implemented as stochastic optical reconstruction microscopy (STORM)(Rust et al. 2006), utilises fluorophore dark-states and sequential imaging to generate lists of molecular coordinates with mean 10 nm precision in fixed cells. Using Bayesian statistical cluster analysis(Griffié et al. 2016; Rubin-Delanchy et al. 2015), it is possible to accurately quantify protein clustering on the nanoscale. This includes clustering of intracellular molecular populations via 3D interferometric SMLM imaging featuring isotropic resolution(Shtengel et al. 2009; Griffié et al. 2017).

We investigated the dynamics of retrograde actin flow and the nanoscale organisation of LFA-1 together with the associated signalling molecules known to be markers of actively signalling adhesions at an early stage in their formation (phosphorylated Src family kinases and FAK) in primary murine T blasts allowed to migrate on coverslips coated with the LFA-1 ligand ICAM-1. A range of conditions were used to modulate cell speed including manganese ions (which push integrins into the high affinity state)(Dransfield et al. 1992), high density ligand (to increase adhesion), the addition of chemokine CXCL12 (to induce signalling through CXCR4) and cytochalasin-D (which inhibits actin polymerisation)(Goddette & Frieden 1986). We included in our analysis cells lacking the phosphatase PTPN22, previously shown to be a negative regulator of migration(Burn et al. 2016). We chose this phosphatase because loss-of-function mutations (Svensson et al. 2011) cause a rapid migration phenotype in T cells and predisposes humans and mice to a range of autoimmune diseases(Dai et al. 2013; Burn et al. 2011; Brand et al. 2005; Bottini et al. 2006; Begovich et al. 2004; Svensson et al. 2011). Using this experimental framework we set out to study the relationship between nanoclustering of LFA-1 and its regulators, and cell speed and actin flow/engagement during polarised migration.

## Results

### The speed of actin retrograde flow is positively correlated with T-cell velocity and negatively correlated with immobile actin

Inside-out and outside-in activation of LFA-1 regulates its affinity maturation and translates into functional effects on adhesion and migration behaviour. F-actin (filamentous actin) forms networks of actin that concurrently ‘slip and grip’(Ponti et al. 2004; Jurado et al. 2005) to facilitate movement by linking highly regulated intracellular flowing actin to the extracellular substrate via transmembrane integrin adhesions(Wiseman et al. 2004; Lawson et al. 2012; Alexandrova et al. 2008; Goult et al. 2013; Changede et al. 2015; Sun et al. 2014). Transient engagement of the actin through adhesions is part of a mechanism that transduces force from flowing actin and is termed the molecular clutch(Chan & Odde 2008; Hu et al. 2007; Brown et al. 2006; Ishibashi et al. 2015; Chen et al. 2012; Case & Waterman 2015; Havrylenko et al. 2014). Flowing actin therefore represents a population that is unengaged with adhesions, whereas immobile actin (in the external reference frame) represents a population in contact with adhesions.

With these features in mind, we first investigated cell speed on a population level (1000s of cells measured) in polarised, migrating murine effector T-cells exposed to a range of stimuli, plated onto ICAM-1 coated glass coverslips. As expected cells slow down when exposed to MnCl_2_ or high surface concentrations of ICAM-1, and speed up in the presence of chemokine CXCL12 as well as in cells in which PTPN22 has been genetically deleted (Supplementary Figure 1). To characterise the dynamics of actin flow in more detail, primary mouse T cells expressing the Lifeact-GFP transgene were allowed to migrate on glass surfaces coated with ICAM-1 before being imaged by TIRF microscopy (Figure 1a). As a proxy for actin linked to the molecular clutch, Fourier analysis was used to determine the fraction of immobile actin in cells, since stationary parts of the actin cytoskeleton must necessarily be engaged with extracellular substrate while the cell crawls over that position. Subsequently, the centre of the cell was continually reset, shifting the frame of reference from the laboratory frame, to the cell frame (Figure 1b). Spatio Temporal Image Correlation Spectroscopy (STICS) was used to measure the velocity and directionality of flowing (unengaged) actin relative to the cell, i.e. from the cell’s frame of reference (Figure 1c).

**Figure 1:**
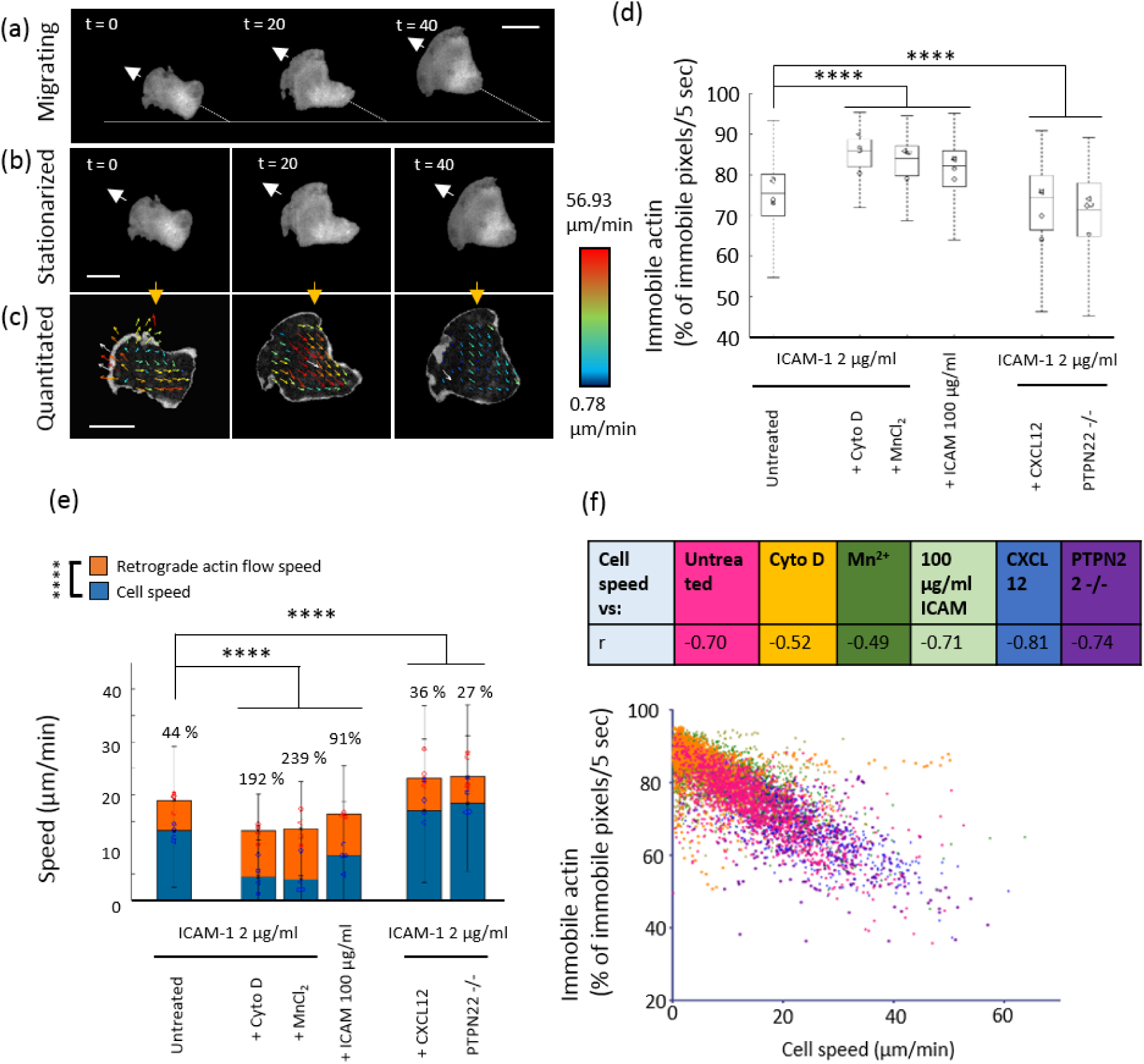
Fast cells exhibit decreased actin attachment to adhesions and increased flow speed. Example frames from representative TIRF movies of actin in a migrating T cell (a). B) stationarised version of the same cell c) STICS vector maps from the stationarised cell. D) Box plot showing immobile actin - % of immobile pixels/5 seconds derived from the external reference frame, E) median cell speed (blue bars – blue symbols are median values for each mouse) and retrograde actin flow speed (orange bars and symbols) from the internal reference frame F) Plot showing negative correlation between % of immobile pixels/5 sec vs cell speed: table above shows r values for correlation in each condition. N = 4 mice, 50 cells per mouse, 200 cells per condition. P<0.00001 ****.

For cells migrating on ICAM-1 (2 µg/ml), 75 ± 5% of pixels contained more than 50% immobile F-actin over 5 second time windows (Figure 1d). Slower cells, treated with cytochalasin D, manganese or a high ICAM-1 concentration (100 µg/ml), displayed a significantly increased percentage of pixels displaying immobile actin p<0.0001 (Figure 1d). In contrast, faster migrating cells (treated with CXCL12 or deficient for PTPN22) had significantly lower percentages of pixels containing immobile actin p<0.0001. Furthermore, Figure 1e shows that in all conditions the fraction of pixels in which actin was immobile was inversely correlated to cell speed (r values for correlation shown in table, p < 0.0001). Therefore, we concluded that decreased actin engagement correlates with increased cell speed.

STICS was then used to analyse the flow speeds of unengaged actin, relative to the cell (Figure 1f). T cells migrating on ICAM-1 (2 μg/ml) alone had an average cell speed of 13.19 μm/min (blue), with retrograde flowing actin moving faster at 18.95 μm/min (orange). Conditions which slowed cell migration also slowed actin retrograde flow speed. Notably, actin flow speed always remained much higher than cell speed in all cases. The percentage values above the bars show that in slower cells, treated with cytochalasin D, Mn2^+^ or a high concentration of ICAM-1, actin speed was 192 %, 239 %, and 91% higher than the cell speed, respectively. Adding CXCL12 to cells or measuring cells deficient for PTPN22 led to faster cell migration which was also coupled to faster actin retrograde flow (Figure 1f). The percentage difference between the speed of actin and the speed of the cell was reduced in comparison to untreated cells and slow cells (36 and 29 %). Taken together, these data indicate that retrograde actin flow is always greater than the overall velocity of the T-cell and that the proportion of immobile actin is inversely related to cell speed.

### The size and number of LFA-1 membrane nanoclusters increase with T-cell speed

When effector T-cells migrate on ICAM-1 coated glass they adopt a polarised morphology (Hogg et al. 2004; Comrie et al. 2015; Valignat et al. 2013; Ridley et al. 2003). Previously, studies of integrin affinity have shown a specific pattern where LFA-1 within the leading edge exists predominantly in the intermediate affinity form, whereas high affinity LFA-1 is detected in the mid cell body/focal zone(Smith et al. 2007). It has been reported in slow moving cells that integrin-based adhesions change their size as they mature(Kanchanawong et al. 2010). We therefore tested whether T cells use differential membrane LFA-1 nano-clustering to regulate cell migration.

Integrin LFA-1 nanoscale clustering in conditions that speed up or slow T cell migration was investigated by dSTORM and Bayesian cluster analysis (Rubin-Delanchy et al. 2015; Griffié et al. 2016). Effector T-cells were allowed to migrate on ICAM-1 coated glass for 15min in the presence of conditioned/unconditioned media, and these were compared with cells deficient for PTPN22. Cells were fixed, surface LFA-1 detected with anti-LFA-1 AF647(clone 2D7), STORM acquisition performed and co-ordinate pointillist maps generated for Bayesian cluster analysis. Stationary effector T-cells were used as non-polarised T-cell controls. Representative STORM images and cluster maps of 2 × 2 µm regions of interest (ROI) within a stationary and migrating T-cell are shown (figure 2a and b). Population cell speeds under each condition were measured by automatic tracking from the same sample, as before (Supplementary Figure 1).

**Figure 2:**
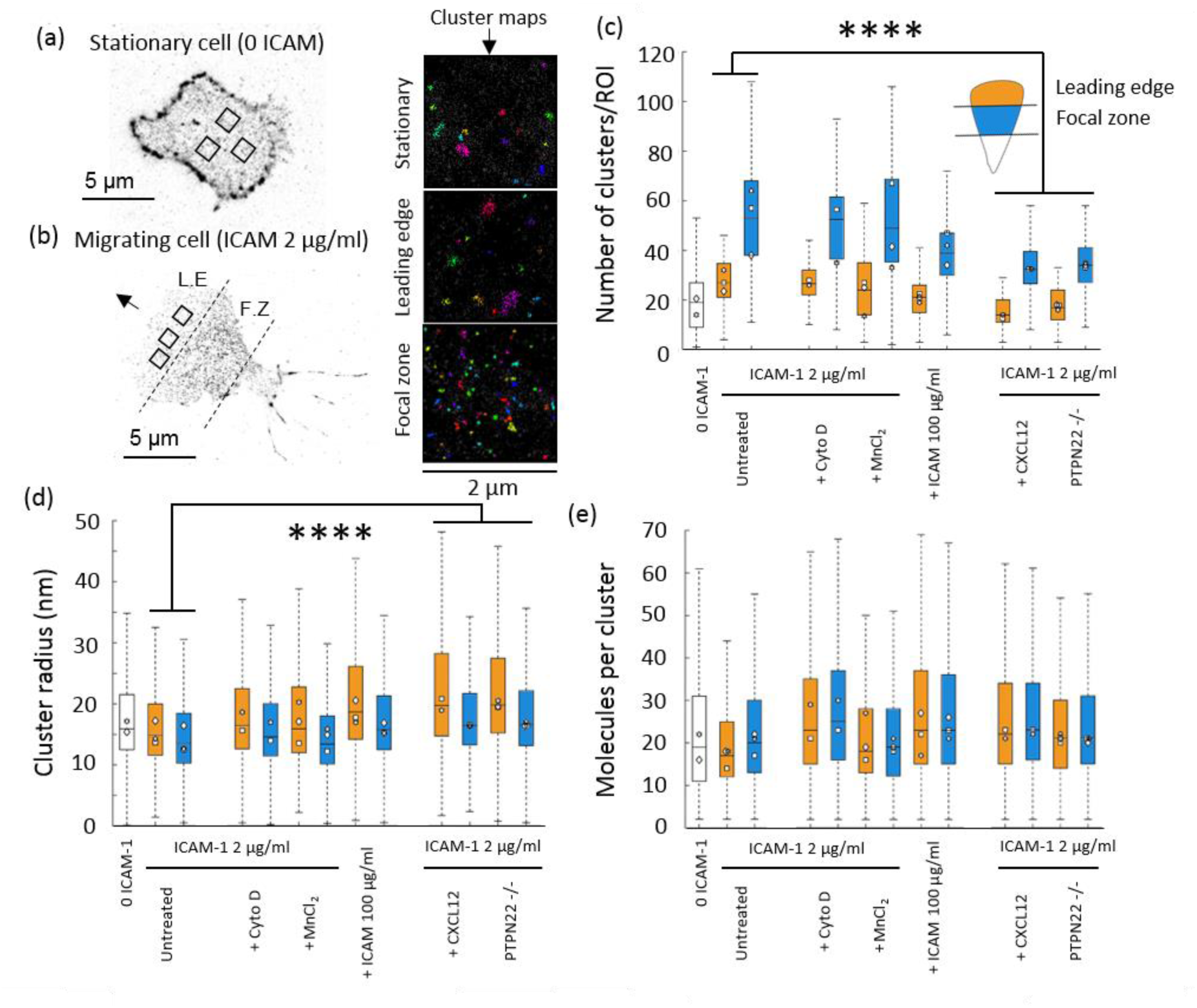
Integrin LFA-1 membrane nanoclusters increase in density and decrease in number in fast migrating cells. Example pointillist maps derived from STORM imaging of LFA-1 in a stationary cells (a) and a migrating cell (b). Boxes in a and b denote 2 μm^2^ regions chosen for analysis and representative cluster maps on the right. In each plot, metrics extracted for stationary cells have clear bars, and migrating cells are split up into leading edge (orange bars) and focal zone (blue bars). From left to right conditions proceed: ICAM-1, 2 μg/ml ICAM-1 coated coverslip with added cytochalasin D, Mn^2+^ or a higher concentration of ICAM-1 (100 μg/ml), then, 2 μg/ml ICAM-1 coated coverslip with CXCL2 added at 150 ng/ml or using cells deficient for PTPN22. Metrics extracted consist of c) number of clusters per ROI, d) cluster radius (nm) and e) the molecules per cluster. (n=350 ROIs and 40 cells per condition from 3 separate mice). P****<0.00001.

In polarised T cells, the focal zone (FZ) displayed different LFA-1 clustering properties to the leading edge (LE). The number of LFA-1 clusters per ROI was consistently increased for example (Figure 2c). The size of clusters was significantly decreased in the FZ (Figure 2d), whereas the number of molecules per cluster was significantly increased (Figure 2e). Together this represents a median increase in cluster density in FZ clusters (blue bars), compared to LE clusters (orange bars). This is achieved both through a size contraction and an increase in molecular content of the clusters. Compared to stationary cells, clusters in the cell membrane of polarised migrating T cells increase in number, in size and in molecular content. Together, this indicates that T cells adopt a regionally discriminated LFA-1 nanoclustering configuration during polarised T cell migration.

The number of clusters significantly decreased in cells that migrated faster due to treatment with CXCL12 or deficiency for PTPN22 (Figure 2c) as compared to slower cells (Mn^2+^ treated, high ICAM-1 concentration and untreated controls). Slower moving cells had similar number of LFA-1 nanoclusters to untreated control cells (Figure 2c). The size of clusters increased in both the LE and FZ of cells treated with CXCL12 or lacking PTPN22 (Figure 2d) however, the number of localisations per cluster was largely unchanged (Figure 2e). Together, this indicates an increase in cluster size, and a decrease in cluster density in fast moving cells.

These results suggest that integrin LFA-1 membrane clusters are modulated on the nanoscale and this influences the migration speed of the cell. Clusters adopt a small dense profile in the high affinity integrin rich focal zone, and a larger less dense profile in the reduced affinity leading edge. Clusters get larger and less dense in faster migrating cells. The data suggest that these zone-specific cluster patterns seem to be hardwired into the polarised T-cell and indicate that the biophysical arrangement of LFA-1 is regulated in conjunction with integrin affinity to control cell speed. As there are fewer clusters in faster moving cells, we next investigated the 3D profile of nanoclusters of LFA-1, to determine whether this phenomenon relates to the recruitment of LFA-1 to the membrane from submembrane stores.

### The size of 3D intracellular LFA-1 clusters increases above the focal zone as cell speed decreases

LFA-1 is constantly recycled to and from the cell membrane to facilitate cell migration. Accordingly, we investigated whether changes in the intracellular pool of membrane proximal LFA-1 was nanoclustered, and if so, was similarly zonally distributed to surface LFA-1 and whether that distribution was modulated with cell migration dynamics. Interferometric photoactivatable light microscopy (iPALM) (Shtengel et al. 2009) was used for the acquisition of 3D single molecule data with ∼20 nm z-precision to a depth of ∼500 nm from the height of the coverslip. The same battery of conditions was used to slow down and speed up migrating T cells, which were tracked to confirm their speed prior to fixation. Figure 3a illustrates a representative iPALM z-projection of cytosolic LFA-1 nanoclusters in a migrating T-cell (left) and the result of Bayesian 3D cluster analysis (Griffié et al. 2017) in a representative 2000 × 2000 × 400 nm ROI (right). Cluster descriptors for the size and number of molecules per cluster were then extracted from ROIs in the FZ or the LE and plotted as a function of z - height from the coverslip.

**Figure 3:**
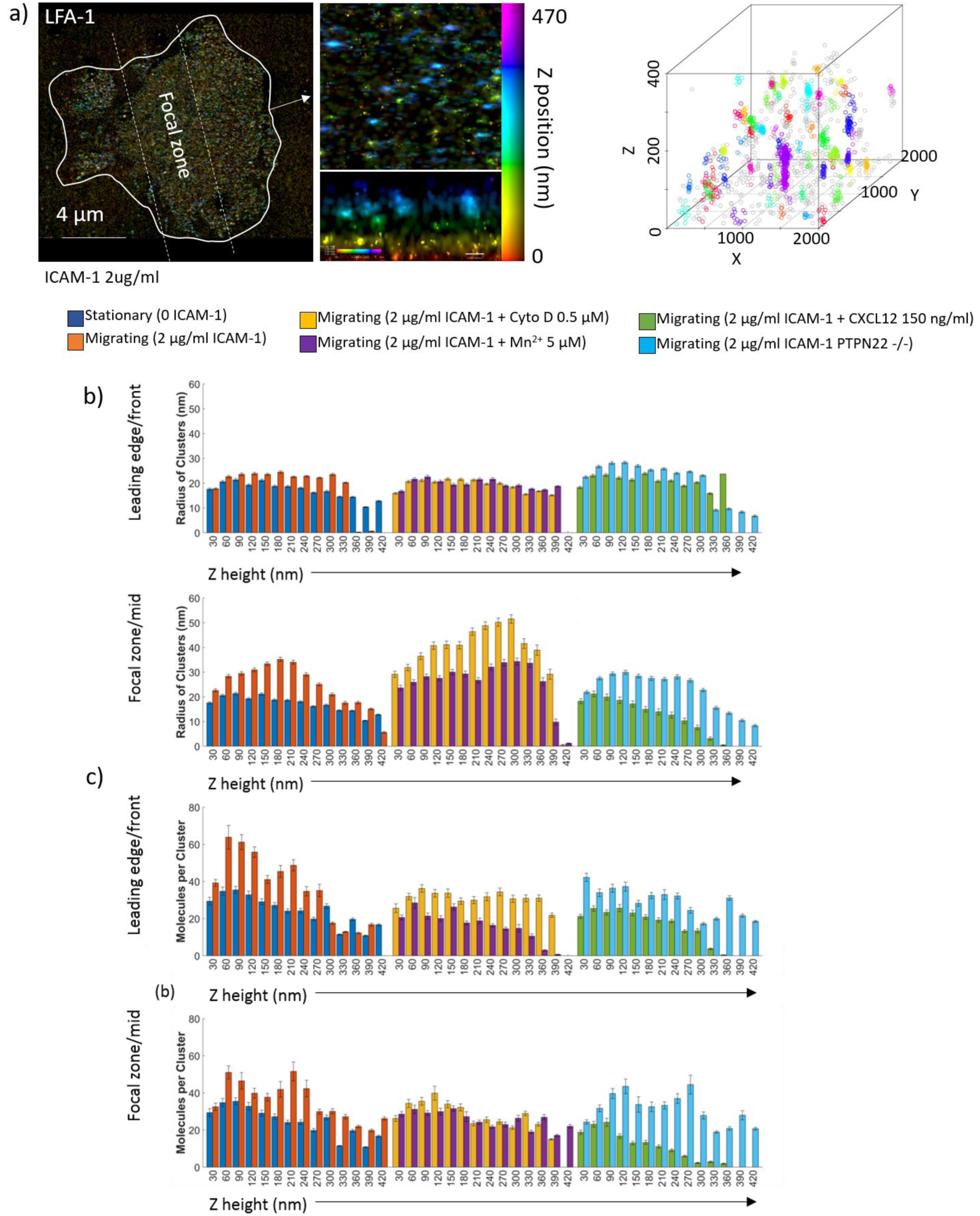
Intracellular LFA-1 nanoclusters are larger in slow moving cells above the focal zone, and smaller in fast moving cells above the focal zone. A) Representative iPalm Z projection of LFA-1 in a migrating T cell (left panel), zoomed region in xy and xz side view (middle panel), and arbitrarily coloured clusters identified in a 2000 × 2000 nm region by Bayesian cluster analysis (right panel). The radius of clusters (b) and the molecules per cluster (c) in were extracted from the cluster maps for stationary cells (0 ICAM-1) and migrating cells (2 µg/ml ICAM-1) (left hand plots), Cytochalasin D and MnCl_2_ treated (middle plots) and CXCL12 and PTPN22 -/- cells (right plots). N = 3 mice, 20 cells and 200 ROIs per condition.

Clustering in migrating cells was regionally distinct between the LE and FZ. In the leading edge, clusters in migrating cells compared to stationary cells exhibited only a small increase in size (Figure 3b – top panel), but this was accompanied by an increase in the number of molecules per cluster (Figure 3c – top panel). Clusters in migrating cells also exhibited an increase in size, focused at a region 200 nm above the coverslip (Figure 3b – lower panel). The number of molecules per cluster was also increased in the FZ of migrating cells (Figure 3c). Together, this indicates that in the switch from stationary to migrating phenotypes, 3D LFA-1 clustering becomes region specific, where clusters above the LE increase in size and density and clusters above the FZ increase in size but maintain the same number of molecules, concentrated close to the cell membrane.

In slowed cells (treated with Cyto D or Mn^2+^), effects on nanoclusters occur above the focal zone, whereas LE clusters remain constant. FZ specific nanoclusters increase in size with a similar number of molecules at all Z-heights in the range 0 to 400 nm. Thus, intracellular LFA-1 nanoclusters become larger and less dense in cells which migrate slower. Conversely in faster cells, a decrease in size of intracellular nanoclusters, with a similar molecular content, was observed above the FZ (light blue and green bars). Deleting PTPN22 (light blue) resulted in a slight reduction in cluster size, however activating T-cells with CXCL12 (turquoise) dramatically reduced the number of LFA-1 molecules per cluster as well as focusing LFA-1 clusters closer to the plasma membrane. Cytosolic pools of LFA-1 are slightly larger in radius, specifically in the FZ compared to membrane.

### Deleting PTPN22 increases T-cell velocity, the size of 3D pY397 FAK nanoclusters and nanoscale colocalization of pY397FAK, pY416Src and LFA-1

Integrin based adhesions are regulated by an array of kinases and phosphatases. Phosphorylation of FAK is a feature of active ICAM-1/LFA-1 signalling (Zhang & Wang 2012; Sanchez-Martin et al. 2004), and is implicated in the first stages of nascent adhesion formation in non-leukocytes (Swaminathan et al. 2016). FAK is basally activated by Src-family kinase LCK in effector T-cells at Y576/577 and Y925. This primes FAK for full activation at the auto-phosphorylation site pY397(Chapman & Houtman 2014) which in turn regulates LFA-1 adhesion and de-adhesion(Raab et al. 2017b). Expression of dominant-negative mutants of FAK in T-cells reduce the speed of migration on ICAM-1 coated glass(Rose et al. 2003), and overexpression of a negative regulator of FAK phosphorylation impairs LFA-1 clustering and reduces ICAM-1/LFA-1 driven T-cell homo-aggregation (Giannoni et al. 2003). PTPN22 is a cytosolic phosphatase that negatively regulates LCK and ZAP-70 downstream of LFA-1 engagement and binds with C-terminal Src kinase (CSK) (Burn et al. 2016), a common target for FAK(Chapman & Houtman 2014). Auto-phosphorylated pY397 FAK has also described as being associated with active integrin in vesicles above the membrane(Nader et al. 2016; Kleinschmidt & Schlaepfer 2017).

We therefore employed iPALM and Bayesian cluster analysis to study 3D pY397 FAK localisation in stationary and migrating cells sufficient or deficient for PTPN22. A representative pseudocoloured 3D localisation map is shown in Figure 4a. ROIs were chosen from the cells, and Bayesian cluster analysis carried out to produce cluster maps (Figure 4b). pY397 FAK staining was verified using FAK inhibitor 14, which blocks Y397 phosphorylation (Supplementary Figure 2). ROIs from the LE or FZ regions were quantified in terms of their radius and molecules per cluster as a function of z (Figure 4c and d).

**Figure 4:**
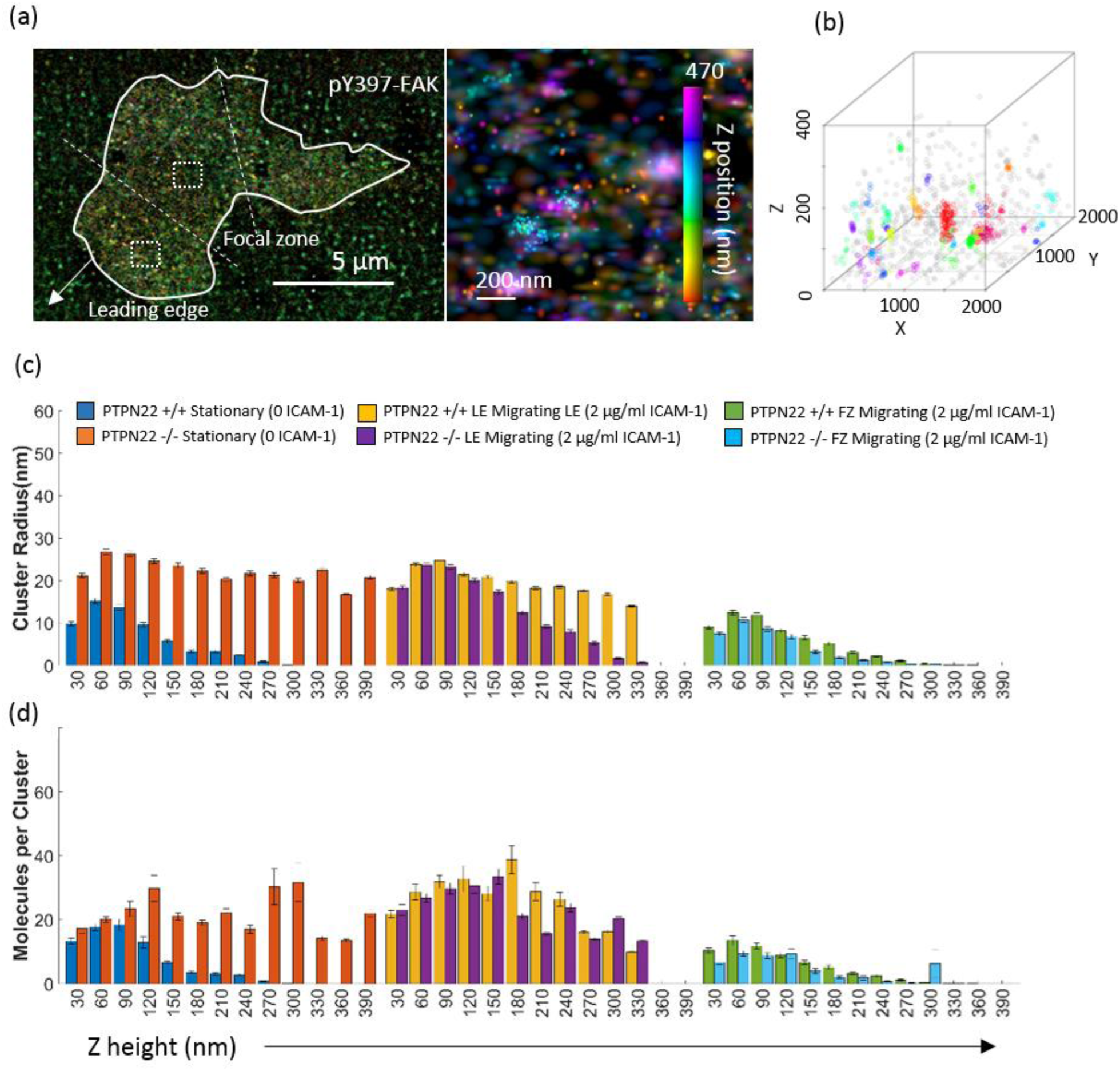
Intracellular 3D pY397 FAK nanoclusters grow larger upon migration, are larger in the leading edge and are preformed high up in stationary cells deficient for PTPN22. a) An example pseudocoloured cluster map (left) is shown alongside a zoomed image where clusters can be visualized at different height scales from 0 to 470 nm (middle panel with z colour bar). b) Shows an example cluster map with defined 3D clusters. For the radius of clusters (c) and the number of molecules per cluster (d), the leftmost plots show stationary PTPN22 +/+ or PTPN22 -/- cells (0 ICAM-1), the middle plots show the leading edge of migrating PTPN22 +/+ PTPN22 -/- cells (2 µg/ml *ICAM*-1) and the right hand plots show the focal zone of migrating PTPN22 +/+ PTPN22 -/- cells (2 µg/ml ICAM-1). N = 3 mice, 20 cells and and 200 ROIs per condition.

When comparing stationary cells to cells migrating on ICAM-1, pY397 FAK cluster size (Figure 4c) increased in the intracellular region above the leading edge but did not increase in the intracellular region above the focal zone. In stationary cells, pY397 FAK nanoclusters were small, and localised close to the membrane (blue bars). Upon ICAM-1 driven cell migration, clusters doubled in size in the LE across a wide z height (yellow bars, 0 to 400 nm), where nascent adhesions were formed after trafficking to the cell membrane (Swaminathan et al. 2016). In the FZ, pY397 FAK clusters remained small (green bars). The molecular content of clusters followed the same patterns as size – larger clusters contained more molecules of pY397 FAK (blue, yellow and green bars figure 4d).

PTPN22 deficient stationary cells (no ICAM-1) adopted larger pY397 FAK nanoclusters (orange bars figure 4c), which remained at this size during cell migration in the LE close to the membrane (purple bars). At points higher in the cell (> 180 nm) clusters were smaller than in PTPN22 sufficient cells (purple compared to yellow bars). Again, clusters above the FZ remained small (light blue bars). The number of molecules per cluster followed the size, becoming greater in larger clusters present in the LE of PTPN22 deficient cells (Figure 4d). Unlike LFA-1 clusters, pY397 FAK clusters maintained their density.

As nanoclusters resemble nascent adhesions in non-leukocytes, but occur throughout the cell, we next investigated how they are regulated in T cells. We tested whether the lack of PTPN22 causes changes in the colocalization of pFAK with LFA-1 and a second set of signalling intermediates: the Src family kinases, using multi-colour SMLM microscopy(Yi et al. 2016a; Yi et al. 2017). Wild type or PTPN22 deficient primary murine T cell blasts were plated on ICAM-1, allowed to migrate and tracked by live-cell imaging. They were then fixed, stained and imaged using madSTORM(Yi et al. 2016b) permitting the sequential imaging of pY397FAK, pY416Src and LFA-1. Supplementary Figure 3 confirmed that PTPN22 deficient cells were faster than PTPN22 proficient cells. Figure 5a shows a representative cell post fixation after three rounds of sequential staining and imaging.

**Figure 5:**
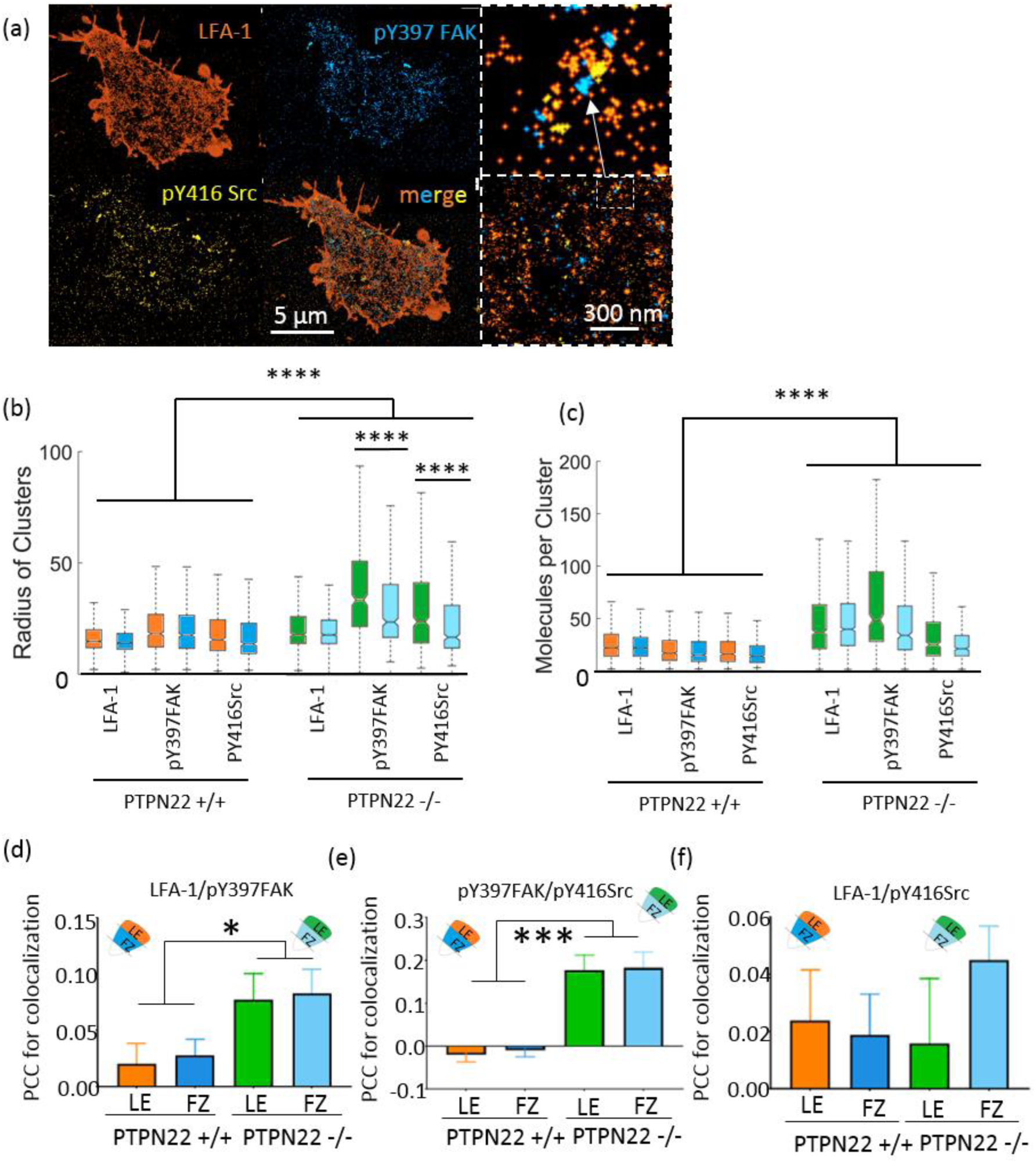
Cluster size and number of molecules per cluster increase in faster moving PTPN22 -/- cells in the case of LFA-1, pY397 FAK and pY416 Src. Individual cells were sequentially stained and imaged by madSTORM: LFA-1. pY397 FAK and pY416 Src in the same cell are shown (a). LFA-1 clusters are larger (b) and denser (c) in faster moving PTPN22 -/- cells. pY397FAK and pY416 Src clusters are also larger and denser in PTPN22 -/- cells. The size of pY397 FAK and pY416 Src clusters in PTPN22 -/- are more strongly regionally discriminated: a) clusters in the leading edge are larger than those in the focal zone, though the molecular content stays the same. Pearson’s correlation coefficient for cluster colocalization: d) LFA-1 colocalized with pY397 FAK more in PTPN22 -/- cells, and pY397 FAK colocalized more with pY416 Src in PTPN22 -/- cells (e). LFA-1 and pY416 Src displayed similar levels of colocalization in PTPN22 +/+ and PTPN22 -/- cells (f). N = 21 PTPN22 +/+ cells and 13 PTPN22 -/- cells. ***p<0.0001.

In wild-type cells, LFA-1, pFAK and pSrc clusters co-existed on a similar size scale and with similar numbers of molecules per cluster (Figure 5b and c). LFA-1, pY397FAK and pY416Src clusters were consistently larger with more molecules in the leading edge compared to the focal zone. PTPN22 deficient cells displayed significantly larger clusters of all three proteins in both the LE and FZ, combined with significantly more molecules per cluster. PTPN22 deficiency therefore increases clustering of all three proteins, but especially pFAK and pSrc. Extraction of Pearson’s Correlation Coefficient (PCC) for the colocalization of clusters of each species showed that LFA-1 clusters were more colocalised with pFAK clusters when PTPN22 was deficient (Figure 5d). pY397 FAK was also more colocalised with pSrc in PTPN22 deficient cells (Figure 5e). The colocalization of LFA-1 clusters with pY416 SFK clusters was unchanged (Figure 5f). Taken together, faster migrating PTPN22 deficient cells display larger LFA-1, pFAK and pSrc clusters which are more often colocalised spatially on the nanoscale.

## Discussion

We employed advanced and super-resolved imaging methods to study the nanoscale migration machinery in T cells. Our data indicate that 1) actin flow and engagement drive migration, where engagement is inversely correlated to cell speed and flow rate of unengaged actin is positively correlated, 2) T cells exploit differential clustering of hundreds of tiny nano-adhesions to translate force from the actin cytoskeleton to the substrate, 3) LFA-1 and pY397 FAK nano-adhesions adopt region specific clustering patterns both in 2D in the membrane and in 3D above the membrane, 4) changes to cell speed induced by distinct experimental conditions modulate the adhesion nano-cluster size and density, and 5) the deletion of PTPN22 phosphatase causes faster T cell migration, and modulates LFA-1 and pY397 FAK clustering in 2D and 3D. Importantly, lack of this phosphatase was sufficient to induce an increase in LFA-1-pYSrc family kinase-pY397FAK nanocluster colocalization coupled to actin flow speed increase, actin engagement decrease and cell migration speed increase.

This is the first time that the front(LE)/middle(FZ) patterns of LFA-1 (2D and 3D), pYFAK (2D and 3D) and pYSrc (2D) have been reported together on the nanoscale. For LFA-1, reported trends for affinity occur in the same zones: high affinity integrin occurs in the FZ, whereas low/intermediate affinity integrin predominates in the LE(Persson et al. 2018; Evans et al. 2011; Smith et al. 2005). In addition, nano-adhesions described here differ from adhesions in slower moving cells, which are founded as ‘nascent adhesions’ and grow as they mature into larger multi-molecular assemblies towards the middle of the cell. Our data suggest a different system, where high affinity LFA-1 adopts smaller, denser more numerous cluster arrays in the FZ. Intracellular LFA-1 clusters are larger above the FZ, and intracellular pY397 FAK clusters are larger and denser above the LE. Coupled with the actin configurations, this indicates that these clusters provide an area of increased adhesion in the FZ: it may be this adhesion imbalance between the LE and FZ that enables fast forward cell migration.

In conditions that increase cell speed, nano-adhesions become larger and are joined by larger clusters of phosphorylated kinases (pY397FAK and pYSrc family). The integration of multiple molecules into the same nanocluster may necessitate its increase in size in this way. That this effect is produced where PTPN22 is knocked-out of cells, which directly targets Lck, a Src family kinase that itself phosphorylates FAK, indicates that the size, density and number of LFA-1 clusters is modulated by such kinases and phosphatases on the nanoscale. Pools of intracellular LFA-1 decrease in density in rapidly migrating cells, and intracellular pY397 FAK clusters stay the same size, indicating a further level of control of adhesion based on the nature of LFA-1 undergoing recycling. *Do we need to make the explicit statement that the changes associated with faster migration make cells less sticky (ie so that they can move faster).*

It is also important to note that T cells are capable of several modes of migration, depending on their environmental niche. Here, we study ICAM-1/LFA-1 dependent migration, coupled to actin treadmilling and engagement. High speed polarised T cell crawling is a phenomenon which occurs in the lymphatic capillaries, during lymph node entry, tissue entry and fast DC scanning. Such integrin dependent migration is especially important during inflammation, and including in autoimmune disease, and is a physiological requirement for high speed migration1. Integrin-free slower migration relies mainly on RhoGTPase mediated actin remodeling and predominates in the dense, fibrous interstitium (ICAM-1 blockade doesn’t affect cell migration here). In the lymph nodes, both integrin independent and integrin dependent fast migration occur, where ICAM-1 blockade reduces T cell migration speed considerably. *In vitro*, the presence of ICAM-1 potentiates migration speed by 50 to 30% in confined 3D (under agarose), although in such a system the cells do not require integrin to migrate slowly. Previously, the simple unconfined ICAM-1 coated glass system used here has been used to characterise integrin affinity states. Here, we have used it to characterise the contribution of nanoclustering and its affect on actin engagement and cell speed.

Overall, we demonstrate that T cells use individually regulated integrin based, kinase/phosphatase coupled ‘nano-adhesions’ to achieve fast migration. Such nano-adhesions are similar to previously described nascent adhesions(Sun et al. 2014) but differ in that they are regulated on a sub-diffraction length scale throughout the cell membrane, and not only in the leading edge. The differential inclusion of proxy markers for integrin activity (phosphorylated FAK and Src family kinases) indicates the highly spatial nature of such adhesion regulation. We posit that this nano-adhesion system, as opposed to a classical nascent > focal complex > focal adhesion system, represents a novel way for T cells to quickly turn over individual adhesions to achieve dynamic feats of migration depending on external or internal cues. The question of live cell dynamics and rare events in the regulation of nanoclusters will be key to understanding how cells use spatiotemporal molecular organisation to change their behaviour. New high throughput techniques(Holden et al. 2014; Gunkel et al. 2014) coupled to low power imaging(Chen et al. 2014), and new fluorescent proteins(Zhang et al. 2016; Chang et al. 2012; Tiwari et al. 2015) may soon enable these kind of investigations, and will be especially important in connecting nanoscale regulation to whole cell function in diverse settings(Shannon & Owen 2019), including in autoimmune disease.

## Methods

### Mice colonies

Male β-actinCre/+; Lifeact-mEGFP/Y mice were were crossed with Ptpn22-/- females; offspring were backcrossed to generate Ptpn22+/+*β-actinCre+Lifeact-mEGFP+* and Ptpn22-/-*β-actinCre+Lifeact-mEGFP+* males. Single cell suspensions of mouse lymph nodes and stimulated with 1 μg/ml of Concanavalin A. After 48hrs T-blasts were re-suspended in medium supplemented with 20ng/ml human recombinant IL-2 for an additional 3-5 days then used in migration assays.

### Live cell phase contrast time lapse microscopy and automatic tracking

was performed using a Nikon Eclipse TI-e microscope. Automatic tracking to quantify T cell migration was performed using a custom ICY modular program. Live cell TIRF time lapse imaging of Lifeact mEGFP+ T cells was performed using a Nikon Ti inverted microscope

### Analysis of actin engagement and flow

Analysis windows or ‘Time of Interest’ (TOI) for actin flow and engagement was 5 frames. Immobile actin was calculated in the external reference frame of the cell by taking the zero component of the temporal Fourier transform. A threshold of 50% was applied to each pixel for it to be counted as immobile. For STICS, we adapted a stationary cell program(Wilson & Theriot 2006), which keeps the cell centroid in the centre of the ROI. Actin flow analysis was then carried out using STICS(Hebert et al. 2005).

### dSTORM

T cells were fixed, blocked and stained with primary directly conjugated antibody. Cells were placed in an oxygen scavenging buffer. Imaging was performed on a Nikon N-STORM microscope. Cells were imaged under TIRF illumination. 10 000 frames were collected at 10 ms exposure time per frame. Molecular coordinates were calculated using ThunderSTORM localisation software(Ovesný et al. 2014) with multi-emitter fitting analysis(Huang et al. 2011) enabled. Cluster analysis was performed using Bayesian analysis in 2D(Rubin-Delanchy et al. 2015) or 3D(Griffié et al. 2017). For iPALM, custom gold nanorod fiducial embedded coverslips(Moore et al. 2018) were coated with ICAM-1. iPALM imaging was performed at the Advanced Imaging Centre at Janelia Research Campus. X, Y and Z coordinates were derived using Peakselector localisation software.

### Single cell correlative live tracking + multicolour dSTORM by multiplexed madSTORM

T cells were first tracked during migration. Cells were fixed, and their positions were recorded, and then multiplexed antibody madSTORM(Yi et al. 2016b; Yi et al. 2017) was used to achieve multicolour imaging. 3 colour pointillist maps were analysed channel per channel with Bayesian cluster analysis(Griffié et al. 2016), and colocalization was analysed(Rossy et al. 2014).

### Statistics

One-way analysis of variance (ANOVA) and Kruskal Wallis post testing was used. All statistical analyses were performed with Graphpad Prism software.

## Acknowledgements

This work was supported by ERC Starter Grant #337187, and Arthritis Research UK Programme grant 20525. We acknowledge use of the Nikon Imaging Centre (NIC), King’s College London. Male β-actinCreER/+; Lifeact-mEGFP/Y mice were a gift from Dr Karen Liu. For the STICS code and support, we acknowledge Paul W. Wiseman and Elvis Pandzic (McGill), with cell stationarity code written by Laurence Yolland and Brian Stramer (King’s College). iPALM imaging was done in collaboration with the Advanced Imaging Center at Janelia Research Campus, a facility jointly supported by the Gordon and Betty Moore Foundation, and the Howard Hughes Medical Institute. PTPN22 KO mice were a kind gift of Rose Zamoyska. Nanodiamonds were a kind gift of Keir Neumann, NIH. M.S. was supported by the King’s Bioscience Institute and the Guy’s and St Thomas’ Charity Prize PhD Programme in Biomedical and Translational Science.

